# Atomistic Characterization of Beta-2-Glycoprotein I Domain V Interaction with Anionic Membranes

**DOI:** 10.1101/2024.03.19.585743

**Authors:** Hale S. Hasdemir, Nicola Pozzi, Emad Tajkhorshid

## Abstract

**Background:** Interaction of beta-2-glycoprotein I (*β*_2_GPI) with anionic membranes is crucial in antiphospholipid syndrome (APS), implicating the role of it’s membrane bind-ing domain, Domain V (DV). The mechanism of DV binding to anionic lipids is not fully understood.

**Objectives:** This study aims to elucidate the mechanism by which DV of *β*_2_GPI binds to anionic membranes.

**Methods:** We utilized molecular dynamics (MD) simulations to investigate the structural basis of anionic lipid recognition by DV. To corroborate the membrane-binding mode identified in the HMMM simulations, we conducted additional simulations using a full membrane model.

**Results:** The study identified critical regions in DV, namely the lysine-rich loop and the hydrophobic loop, essential for membrane association via electrostatic and hydrophobic interactions, respectively. A novel lysine pair contributing to membrane binding was also discovered, providing new insights into *β*_2_GPI’s membrane interaction. Simulations revealed two distinct binding modes of DV to the membrane, with mode 1 characterized by the insertion of the hydrophobic loop into the lipid bilayer, suggesting a dominant mechanism for membrane association. This interaction is pivotal for the pathogenesis of APS, as it facilitates the recognition of *β*_2_GPI by antiphospholipid antibodies.

**Conclusion:** The study advances our understanding of the molecular interactions be-tween *β*_2_GPI’s DV and anionic membranes, crucial for APS pathogenesis. It highlights the importance of specific regions in DV for membrane binding and reveals a predominant binding mode. These findings have significant implications for APS diagnostics and therapeutics, offering a deeper insight into the molecular basis of the syndrome.

## Introduction

Beta-2-glycoprotein I (*β*_2_GPI), also known as apolipoprotein H, is a heavily glycosylated protein present in the blood.^1–5^ Despite its abundance and involvement in complement and clotting cascades, its exact physiological function remains unclear. Nevertheless, it is well-established that it plays a crucial role in antiphospholipid syndrome (APS) as it is the primary target of antiphospholipid antibodies (aPLs).^6–11^ In APS, *β*_2_GPI recognition by aPLs is believed to trigger the activation of several cell types, most notably platelets, monocytes, neutrophils, and vascular endothelial cells.^9^ This activation then leads to dysregulation of complement and coagulation pathways, resulting in inflammation, vasculopathy, thrombosis, and pregnancy-related complications, all of which have been documented in APS patients.^9,12–14^

An important feature of APS is that pathogenic aPLs preferentially recognize *β*_2_GPI when bound to anionic phospholipids such as phosphatidylserine (PS) and cardiolipin (CL). These anionic phospholipids are usually located inside healthy resting cells but become exposed to the extracellular environment upon cell activation, autophagy, or necrosis, thus explaining why underlying inflammatory conditions, infections, or similar clinical scenarios may trigger thrombosis in APS patients.^11,15^ Consequently, binding of *β*_2_GPI to anionic phospholipids is essential for APS pathogenesis.^9,16^

*β*_2_GPI is comprised of 326 amino acids arranged in four canonical complement control protein (CCP) domains (domains I-IV), and one aberrant CCP domain (domain V; DV) (Fig. 1).^2,3,5^ DV is the primary domain for *β*_2_GPI’s interaction with anionic phospholipids, thus playing a pivotal role in the membrane-binding process. ^9,16–21^ Emerging research suggests that although pathogenic aPLs can interact with the freely circulating form of *β*_2_GPI, membrane-binding of *β*_2_GPI through DV significantly enhances antibody recognition possibly by localizing *β*_2_GPI on the cell surface and thus increasing the concentration of the antigen available for recognition.^20–22^ This study aims to investigate the mechanism by which DV binds to membranes and interacts with anionic lipids at an atomic level, providing new insights into its pivotal role in APS.

**Figure 1:**
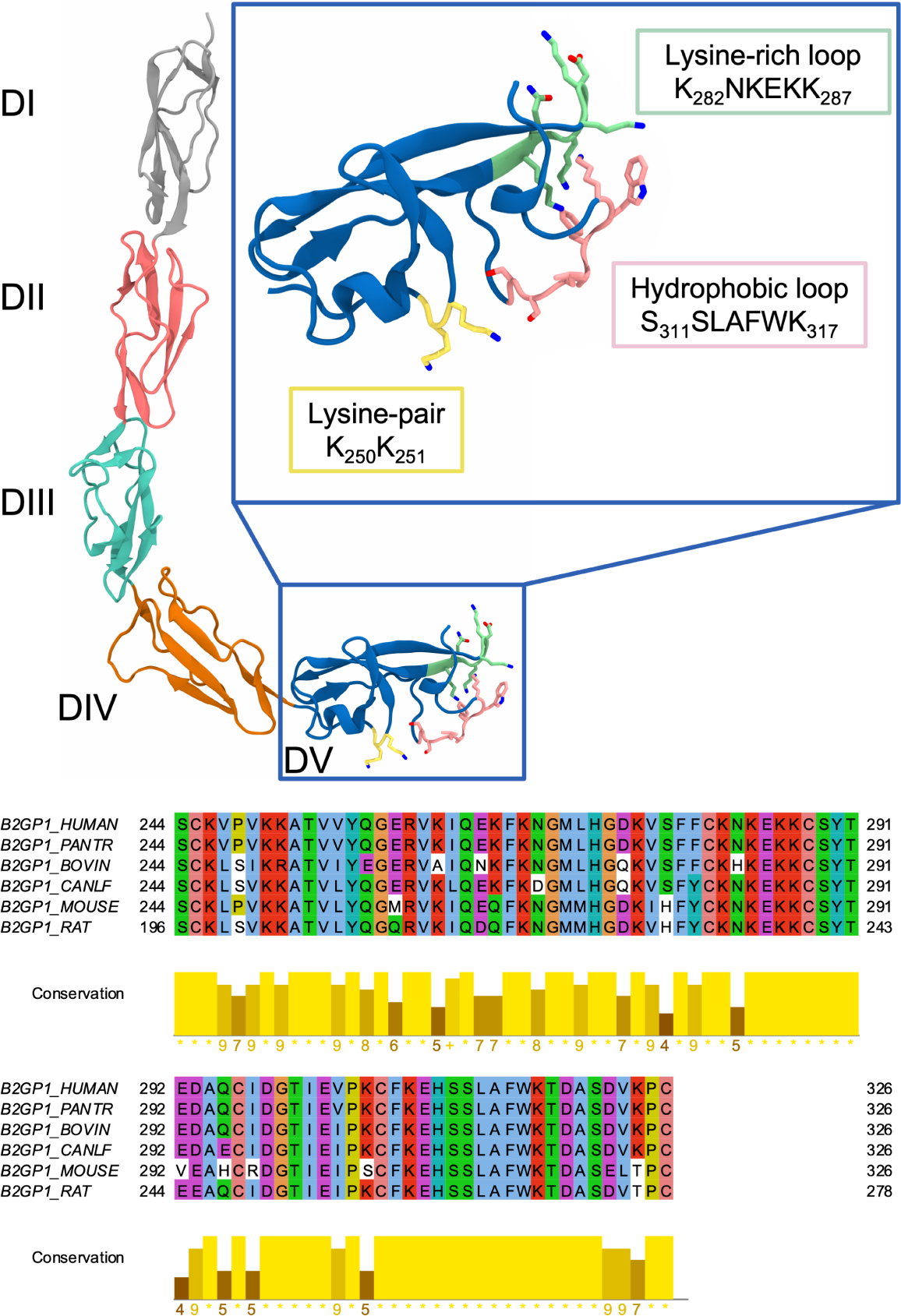
Structural overview of *β*_2_GPI and multiple sequence alignment of mammalian DVs. Top: X-ray crystal structure showcasing the J-elongated conformation of *β*_2_GPI. Domains DI-DV are depicted in gray, pink, green, orange, and blue, respectively. The inset offers a close-up view of DV, with important structural features, namely, the lysine pair, the lysine-rich loop, and the C-terminal, hydrophobic loop highlighted in yellow, green, and pink, respectively. Bottom: multiple sequence alignment of DV from six mammalian *β*_2_GPI proteins. The sequences from top to bottom are: human, chimpanzee, bovine, dog, mouse, and rat. Each column is color-coded based on the type of the most conserved residue at that position. Hydrophobic (A, I, L, M, F, W, V), aromatic (H, Y), polar (N, Q, S, T), acidic (E, D), and basic (K, R) residues are represented in blue, cyan, green, magenta, and red, respectively. Cysteines, prolines, and glycines are shown in pink, yellow, and orange, respectively. Not conserved residues are presented in white. Below the alignment, a conservation score, denoting the number of conserved physicochemical properties for each column, is provided. Strictly conserved residues are marked with ‘*’, while columns with mutations conserving all physicochemical properties are indicated with ‘+’. Columns with higher sequence variance are labeled with the corresponding numerical score.

Domain V (DV) features a largely electropositive region, characterized by multiple conserved lysine residues (K282-K287), near its C-terminal hydrophobic loop (S311-K317) (Fig. 1).^2,3,5^ Several studies have proposed that this lysine-rich loop (K282-K287) in DV can directly interact with the negatively charged headgroups of anionic phospholipids, ^23–30^ while the flexible hydrophobic loop containing W316 is hypothesized to insert into the membrane.^17,26,26–33^ A blue shift in the intrinsic fluorescence spectrum of *β*_2_GPI in the presence of liposomes, which mostly reports on the chemical environment sensed by W316, is consistent with this model.^17,29,34^ Yet despite these observations and considerable sequence conservation of DV across mammals (Fig. 1), the exact role of these and other residues remains incompletely understood.

Investigating the membrane-binding mechanism of a peripheral protein like *β*_2_GPI and its interactions with particular lipids at an atomic detail presents considerable challenges. Computational methods, particularly molecular dynamics (MD) simulations, are an ideal approach for characterizing the intricacies of these biologically relevant processes. In a previous study, interactions between *β*_2_GPI and neutral membranes were investigated using MD simulations, focusing on the primary phospholipids found in synovial fluid, i.e., 1,2-dipalmitoyl-sn-glycero-3-phosphocholine (DPPC) and 1-palmitoyl-2-oleoyl-sn-glycero-3-phosphoethanolamine (POPE).^35^ Notably, both DPPC and POPE are neutral lipids, and the initial positioning of the protein was on the membrane surface. Here, we leverage a type of membrane model called the highly mobile membrane mimetic (HMMM) to investigate the spontaneous membrane-binding process. The HMMM model is a powerful technique that overcomes the limitations of traditional MD simulations for complex membrane systems, which include slow-dynamic lipid molecules in them. The model is particularly tailored for studying protein-lipid interactions of peripheral membrane proteins.^36–41^ The HMMM model has demonstrated utility across a spectrum of membrane-associated phenomena, including membrane binding of cytochrome P450, ^42^ the PH domain,^43,44^ the GLA domains of coagulation proteins,^45–47^ the SH2 domain,^48^ and the SARS-CoV2 fusion peptide,^49^ where it has provided microscopic insights into the membrane-binding mechanism of these proteins.

Using the HMMM model, we identified the structural basis of membrane recognition by DV, providing for the first time an atomic view of how this domain approaches, binds, and interacts with the anionic membranes. Our findings highlight the critical role played by the positively charged lysine-rich loop in the surface-level interaction of DV with anionic lipid headgroups, as well as the anchoring of the hydrophobic loop in the core of the membrane. Additionally, our simulations unveiled a previously unreported membrane interaction site– a lysine pair (Fig. 1)–providing novel insights into the current understanding of *β*_2_GPI membrane binding. These results greatly advance our understanding of membrane binding of *β*_2_GPI, which is key for APS pathogenesis.

## Materials and Methods

### Multiple sequence alignment

Multiple sequence alignment of DV across various mammalian species was performed to identify conserved residues and putative functional hotspots. The alignment included six known mammalian sequences from human, chimpanzee, bovine, dog, mouse, and rat. The alignment of these six sequences was conducted and visual-ized using `Jalview`, utilizing the default `ClustalW` settings and physicochemical proper-ties^50–52^(Fig. 1).

### Protein preparation

The structure of DV was obtained from the human *β*_2_GPI crystal structure purified under physiological pH and salt conditions ^5^ (PDB: 6V06). DV was extracted from the rest of the protein in VMD (Visual Molecular Dynamics).^53^ DV was capped with an acetylated amino terminus (ACE), and disulfide bonds were introduced at C245-C296, C281-C306, and C288-C326 using the `PSFGEN` plugin of VMD. The protonation states of protein residues were determined by pKa calculations using the Protonate-3D module in MOE (Molecular Operating Environment, 2019.01; Chemical Computing Group ULC). The final DV protein structure was then used to generate protein-membrane systems using `CHARMM-GUI`.^54^

### Membrane binding simulation setup

A major challenge in using conventional MD simulations to capture membrane binding is sufficient sampling of potential binding poses. The characteristically slow lateral diffusion of membrane lipids frequently results in suboptimal sampling of protein configurations on the surface of the membrane within the timescales achievable by atomistic MD simulations. ^55^ Consequently, the outcomes can be skewed, reflecting biases introduced by the initial lipid distribution and protein placement. Addressing this limitation, the HMMM membrane model enhances the lateral diffusion of lipids by using short-tailed lipids and employing a membrane core filled with a hydrophobic liquid. ^39,40^ This technique effectively bypasses some of the limitations inherent in traditional MD simulations, enabling increased sampling while retaining atomistic details of lipid-protein interactions on the surface of the membrane. ^39,40^ In this study, we utilized HMMM membranes to initially capture the spontaneous membrane binding of DV. The procedure for constructing protein-membrane systems is outlined in detail below.

To create protein-membrane systems, first 90 × 90 × 46 Å^3^ full membrane systems (including *∼*100 lipids in each leaflet) with different lipid compositions were constructed using `CHARMM-GUI Bilayer Builder`.^54,56^ Then DV was placed in solution with its center of mass (COM) 10 Å away from the membrane surface. The lipid compositions with anionic lipids used (Table 1) were symmetric, as in apoptotic cell membranes the lipid asymmetry between the inner and outer leaflets is lost and negatively charged phospholipids are also exposed to their outer surfaces.^57^ The exposed negatively charged lipids are subsequently recognized by the *β*_2_GPI.^16–19,58^ Additionally, a control case lacking negatively charged phospholipids was included in the set for comparison (Table 1).

**Table 1:** Details of the HMMM simulation systems with varying concentrations of phosphatidylcholine (PC) and PS lipids and starting with different orientations of DV in solution. All simulations were conducted for at least 300 ns. Slight variations in simulation times were observed due to variable termination of MD simulations on supercomputers.

| Membrane Composition | DV orientations | Replicas | Time (ns) per replica |
| --- | --- | --- | --- |
| PC:PS (50:50) | 5 | 3 | $\geq 300$ |
| PC:PS (80:20) | 5 | 3 | $\geq 300$ |
| PC:PS (100:0) | 5 | 3 | $\geq 300$ |

Then, full membrane systems were converted to HMMM membranes by removing atoms after C26 and C36 (the sixth carbon atoms) in the two phospholipid acyl chains. An organic solvent, named Simple Carbon Solvent Ethane (SCSE)^40^ that consists of two carbon-like particles, was used to fill the membrane core, matching the number of carbon (CH2 or CH3) atoms removed in the previous step. To achieve better sampling of protein-membrane interactions, the protein’s initial orientation was randomized by rotating around *x* and *y* axes. The initial position of protein, obtained from `CHARMM-GUI`, was considered as the reference and named (0,0). The protein was then rotated *±*45 deg around *x* and *y* axes independently, to create four additional different starting orientations, which are referred to by the degree of rotations around the two axes, namely (45, 0), (−45, 0), (0, 45), and (0*, −*45) (Fig. S1A). Then, three simulation replicas were generated for each orientation, resulting a total of 15 simulation systems per lipid composition. The systems were solvated using the `SOLVATE` plugin and neutralized with 0.15 M NaCl using the `AUTOIONIZE` plugin of VMD. The final simulation systems consisted of 73,000-82,000 atoms.

### Conversion of HMMM-bound DVs to full membranes

To ensure the stability of the membrane-bound conformations captured by the HMMM model, a representative frame– specifically, the medoid structure of the PC:PS (50:50) membrane composition from cluster 1 (see Analysis below)–was converted from HMMM to full membrane. This conversion involved the removal of the SCSE solvent molecules and the elongation of short-tailed lipids, achieved using the `PSFGEN` plugin in VMD. The resulting full-membrane (FM) system was minimized for 10,000 steps and subsequently equilibrated in six steps under the conditions outlined in Table 2. During this process, the C*_α_* atoms of DV were harmonically restrained. Additionally, the COM of the head group heavy atoms of POPC and POPS lipids were restrained along the *z* axis (membrane normal) to +19 or −19 Å (with membrane midplane at *z* = 0) in the upper and lower leaflets, respectively, using the colvars module of NAMD.^59^ This stepwise equilibration protocol ensured the melting of newly elongated lipid tails. Then, the whole system was simulated for an additional 300 ns without any restraints.

**Table 2:** Details of the equilibration protocol of the DV full-membrane system.

| Equilibration Step | Ensemble | Force Constant on Protein (kcal/mol/Å<sup>2</sup>) | Force Constant on Lipids (kcal/mol/Å<sup>2</sup>) | Time (ps) |
| --- | --- | --- | --- | --- |
| 1 | NVT | 10.0 | 5.0 | 250 |
| 2 | NVT | 5.0 | 2.0 | 250 |
| 3 | NVT | 5.0 | 1.0 | 250 |
| 4 | NPT | 5.0 | 1.0 | 500 |
| 5 | NPT | 1.0 | 0.2 | 500 |
| 6 | NPT | 0.1 | 0.0 | 500 |

### Simulation protocol

The protein-HMMM simulations were performed using NAMD 2.14^60^ and the protein-full membrane simulation with NAMD 3.0.^61^ A timestep of 2 fs was used for both. CHARMM36m^62^ and CHARMM36 parameters^63^ were used for protein and lipids, respectively. TIP3P model was used for water.^64^ The system was first energy-minimized for 10,000 steps using steepest descent algorithm followed by equilibration MD for 5 ns. In the equilibration phase, C*_α_* atoms of the protein and C2, C26, and C36 atoms of lipids were restrained with a force constant of 1 kcal/mol/Å^2^. The temperature was maintained at 310 K using Langevin thermostat (with a damping coefficient, *γ* = 1 ps^−1^), and the pressure was maintained at 1 bar using the Nośe-hoover piston method.^65,66^ The particle mesh Ewald (PME) method was used to calculate electrostatic interactions ^67^ without truncation within the periodic boundary conditions in all the simulations. The non-bonded forces were calculated at 12 Å cutoff with 10-Å switching distances. To contain SCSE molecules in the core of the HMMM membranes, a grid-based potential was applied using the `gridForce`^68^ feature of NAMD. The protein-membrane systems were designed with a flexible cell that permitted independent adjustments in three dimensions for the system size, while maintaining a constant *x/y* ratio of the membrane. Bonds with hydrogens were maintained rigid using SHAKE^69^ and SETTLE algorithms.^70^ The production simulations were carried out for at least 300 ns for each replica, collecting an aggregate of 13.5 *µ*s sampling of DV interaction with membranes.

### Analysis

The number of contacts (C) between the protein and membrane was calculated using the following equation:

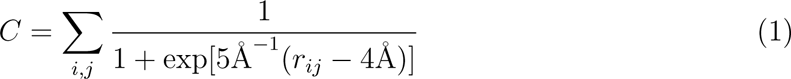

Here, the summation encompasses all possible heavy atom pairs between the protein and lipids, with *r_ij_* representing the inter-atomic distance in a specific frame of the MD trajectory. C can be defined for individual residues, domains, or the entire protein. Similar contact functions have been used previous simulation studies to quantify molecular-level interactions.^71,72^

To pinpoint the membrane-interacting hotspots on DV, we utilized Equation 1 to compute the number of contacts between each residue and different membrane components (Fig. S1B). Subsequently, this quantity was normalized based on the total number of frames in which C*>*0.

To assess the potential preference of DV for anionic phospholipids, we computed C between the protein and PC as well as PS lipids in systems featuring a membrane composition of PC:PS (50:50) (Fig. 3A). Subsequently, the sum of these contacts was normalized by the number of frames in which C*>*0 for the PC:PS (50:50) systems.

To quantify the interaction between various segments of PS lipids and DV, the heavy atoms of PS lipids were grouped into six groups: serine carboxy, serine amine, serine backbone, phosphate, glycerol, and tails (Fig. 3B). In systems featuring the membrane compositions PC:PS (50:50) and PC:PS (80:20), C was computed between each residue and each segment of the PS lipids across the trajectory (Fig. 3C). For each membrane composition, the cumulative C values for each residue were then normalized by the number of frames with C*>*0.

In order to differentiate surface-level membrane interactions from membrane insertion, we measured the *z*-distance between the phosphate plane in the cis leaflet (the leaflet interacting with the protein) and the average side-chain COM of a specified part of the protein. The phosphate and amino planes were defined as the average *z* coordinates of phosphorus and nitrogen atoms, respectively, throughout the simulation. Furthermore, the *z*-distances of the hydrophobic loop, the lysine-rich loop, and the lysine pair were determined with respect to the phosphate plane using the average *z*-coordinate of their side-chain COM. Negative *z*-distances signify membrane insertion.

Using the contact metric (C), frames were categorized as “membrane-bound” if C *≥* 10 (Fig. S6). To exclude transient interactions, only segments of the simulations, where DV remained bound to the membrane for at least 50 ns, were considered for further analysis.

For each “membrane-bound” frame, individual residue distances were computed as the *z*-distance between the phosphate plane and the side-chain COM of each residue (Fig. S6B). The data from these frames were standardized to unit variance and zero mean using the scikit-learn package.^73^ The resulting dataframe included 5,979 rows representing the total number of membrane-bound frames from PC:PS (50:50) and PC:PS (80:20) simulations, and 83 columns, representing the number of DV residues. Dimensionality reduction was performed using Uniform Manifold Approximation and Projection (UMAP) v.0.3.2,^74^ reducing the number of columns from 83 to 2. UMAP maintains proximity of similar data points in the high-dimensional space in the reduced-dimensional space while preserving the global structure of the dataset. UMAP has been successfully employed for identifying and clustering similar biological datasets into lower-dimensional spaces.^75–77^ To achieve well-separated clusters, UMAP parameters were set as follows: n neighbors = 30, min distance = 0.1, n components = 2, random state = 100, Euclidean distance metric. Subsequently, DBSCAN (Density-Based Spatial Clustering of Applications with Noise)^78,79^ was applied to the two UMAP dimensions to classify membrane-bound conformations. DBSCAN, a density-based clustering method, identifies clusters with varying shapes and sizes. Parameters were set to: min samples = 100, and eps = 1. Outliers detected by DBSCAN were excluded from the clustering results. The clusters of membrane-bound states were then projected onto the two UMAP dimensions. For each membrane composition, the frame closest to the cluster average was selected as the representative membrane-bound frame of that cluster. The determination of the representative frame involved calculating two medoids, one for each membrane composition, within each cluster. A medoid was defined as the frame with the lowest total pairwise distance to the cluster average of the original dataframe (5,979 rows and 83 columns).

To characterize the orientation of membrane-bound DV configurations, we evaluated the angle between the three principal axes of DV and the membrane normal at each frame within each cluster. This analysis of the distribution of principal axis angles allowed us to observe the variability in the rotational degrees of freedom for each membrane-bound conformation, providing insights into the stability of DV conformations when bound to the membrane. The analysis was conducted using VMD and the Python packages Matplotlib, Seaborn, Scikit-learn, and SciPy.^73,80–83^

## Results and Discussion

*β*_2_GPI is recognized with high affinity by aPLs only upon membrane association. Consequently, its interaction with anionic phospholipids is pivotal in precipitating thromboembolic events in APS patients. Nevertheless, the details of *β*_2_GPI’s interaction with anionic phospholipids remain elusive. In this study, we employed atomistic MD simulations to describe the spontaneous membrane binding of DV and characterize its associated membrane binding modes. Firstly, we describe DV hotspots that drive spontaneous membrane association by quantifying protein-lipid contacts and membrane insertion. Subsequently, we determine different membrane binding modes of DV by analyzing the relative position of each residue with respect the membrane and clustering them. Further, we assess the stability of membrane binding modes by measuring the scatter of angles between the principal axes of the DV and the membrane normal. The physiological relevance of our membrane-bound configuration is then examined by showing that it is consistent with binding of full *β*_2_GPI structure to the membrane.

### Spontaneous membrane binding of DV

To capture the spontaneous membrane binding of DV, we performed 45 independent MD simulations of DV in the presence of membranes containing PC and PS in varying compositions (Table 1). Membrane compositions comprising 20% and 50% PS were used to mimic apoptotic cell membranes, which expose anionic lipids to the extracellular side. A membrane composition of 100% PC was used as control. We employed the HMMM membrane model to enhance lipid dynamics and mixing, which are known to slow down membrane-associated phenomena in simulations.^55^ The protein was placed initially in five distinct orientations with respect to the membrane surface (Table 1, Fig. S1A) to ameliorate the initial bias in sampling membrane-bound protein forms. To quantify the specific regions on DV that interacted with the membrane, we calculated the number of contacts (C) between each residue and the membrane (see Methods). The simulations highlight three primary membrane interaction sites on DV: the hydrophobic loop (S311-K317), the lysine-rich loop (K282-K287), and the lysine pair (K250/K251) (Fig. S1B). We observe an increase in the number of contacts between two positively charged regions of DV (the lysine pair and the lysine-rich loop) in the context of 50% PS composition compared to 20% PS composition. In contrast, the hydrophobic loop displayed numerous interactions with PS-containing membranes regardless of PS concentration. These results show that both electrostatic forces and hydrophobic effects drive DV and the membrane together. Notably, minimal to no interaction between DV and the membrane occurred in the absence of PS. This observation highlights the critical role of anionic lipids, in this study PS, as a pivotal factor in facilitating the interaction of DV with the plasma membrane. This discovery aligns with previously reported experimental outcomes, such as the absence of membrane binding of B2GP1 to various *in vitro* membrane models composed solely of neutral lipids such as PC, sphingomyelin, and phosphatidylethanolamine.^16,29,34,84–90^

To differentiate DV hotspots that contribute to surface-level interactions with lipids from those anchoring DV in the membrane, we measured the *z*-distance between the bilayer phos-phate plane and the side chains in the hydrophobic loop, the lysine-rich loop, the lysine pair, and other selected sites (Figs. 2 and Figs. S2 to S5). Our results reveal that in the membrane-bound DV, the hydrophobic loop penetrates beneath the phosphate plane in membranes containing PS, indicating a potential anchoring mechanism for DV to the membrane (Figs. 2 and Figs. S2 and S3). The average insertion times for the hydrophobic loop during the simulations were 123 ns and 161 ns for membrane compositions with 50% and 20% PS, respectively (Fig. 2). Moreover, in the majority of our simulations, the lysine-rich loop and the lysine pair consistently contributed to membrane binding through surface-level, electrostatic interactions with anionic lipids (Figs. S2, S4 and S5). Previous experimental studies indicated a decreased binding affinity of DV to anionic membranes when the concentration of Na^+^ or Ca^2+^ in the buffer was increased, which potentially shields the negatively charged membrane surface.^86,88,91^ Our findings suggest a dynamic interplay between peripheral interactions and membrane insertion during DV binding to the membrane, with distinct roles attributed to the hydrophobic loop, the lysine-rich loop, and the lysine pair.

**Figure 2:**
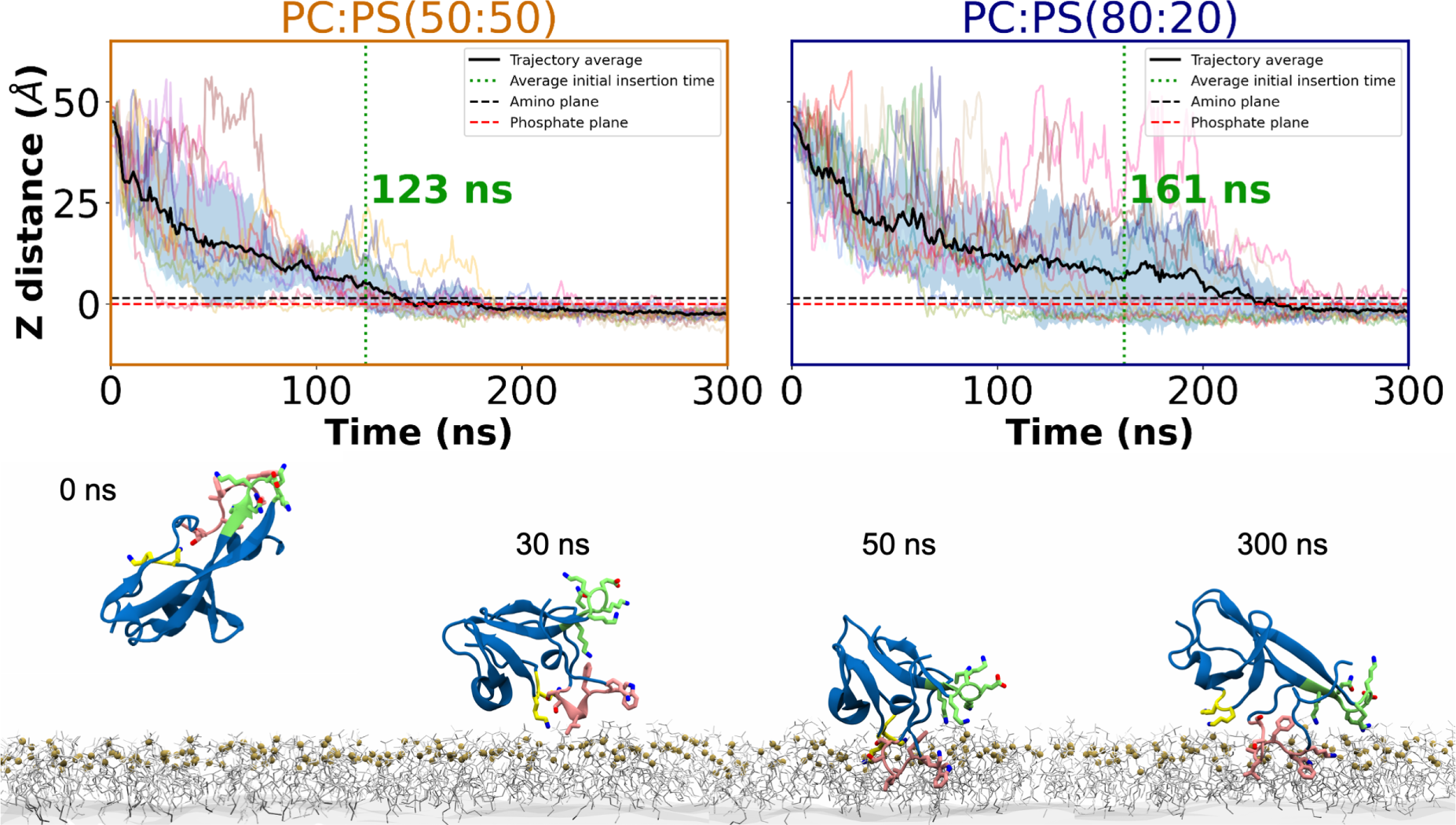
Spontaneous membrane binding of DV via the insertion of the hydrophobic loop into the membrane. Top: time evolution of *z*-distance between the side chain COM of the hydrophobic loop relative to the phosphate and amino planes for selected replicas exhibiting membrane insertion. Light-colored lines show the individual time series, whereas the black line shows the average of all replicas, with the standard deviation shown as a blue region. DV is considered to be inserted in the membrane when *z*-distance between the hydrophobic loop and phosphate plane is negative. The positions of the phosphate and amino planes of the cis-side lipids are indicated using red and black dashed line, respectively. The green dashed vertical line marks the average time for observing membrane insertion. Bottom: Representative snapshots at different time points of a representative simulation illustrating the steps of DV binding to the membrane accompanied by the anchoring of the hydrophobic loop. DV is presented mostly in blue, with the lysine pair, the lysine-rich loop, and the hydrophobic loop highlighted in yellow, green, and pink, respectively. Phospholipids are shown in gray, and the organic solvent in the membrane core is represented as gray surface. The phosphorous atoms of lipids are highlighted as tan spheres.

Previous studies have established the necessity of anionic lipids, specifically PS and car-diolipin, for the membrane binding of both *β*_2_GPI and isolated DV.^16,86,88–90^ To determine DV’s lipid selectivity, specifically between anionic (PS) and neutral (PC) lipids, we assessed the contacts between DV and each lipid type in PC:PS (50:50) simulations (Fig. 3). Our results show that the lysine-rich loop and the lysine pair predominantly interact with the PS headgroups (Fig. 3A). Additionally, the hydrophobic loop exhibits twice as much in-teraction with PS lipids compared to PC lipids (Fig. 3A). We attribute this observation to the protein-lipid closer interaction primed by the anionic lipids. Further analysis of the interaction between DV and PS lipids involves dividing the PS lipids into six sub-domains (Fig. 3B) and quantifying the contacts between each sub-domain and DV residues individu-ally (Fig. 3C). The lysine pair and the lysine-rich loop primarily interact with the negatively charged serine carboxy group and, to a lesser extent, with the phosphate group of PS lipids. This aligns with our previous results, indicating surface-level interactions between these residues and the membrane (Figs. S2, S4 and S5 and Fig. 3C). K308, near the hydrophobic loop, and K317 within the hydrophobic loop interact with the serine carboxy groups, while K317 engages also with the phosphate groups. Our results are validated by previous findings indicating that point mutations at K308, the lysine-rich loop and the hydrophobic loop sig-nificantly diminish the binding affinity of DV to anionic lipids, specifically to cardiolipin. ^92^ K262 and K268 also contribute to surface-level interactions with serine carboxy groups to a lesser degree compared to the aforementioned positively charged residues. L313, A314, F315 and W316 in the hydrophobic loop insert into the membrane hydrophobic core and interact with lipid tails (Fig. 2 and Figs. S2 and S3). Notably, the pronounced interactions observed between the PS lipid tail region and W316 emphasize the critical role of this residue in membrane binding. This observation supports previous experimental findings indicating the membrane anchoring of W316 and demonstrating loss of *β*_2_GPI-membrane binding in W316S mutants.^17,27,34^

**Figure 3:**
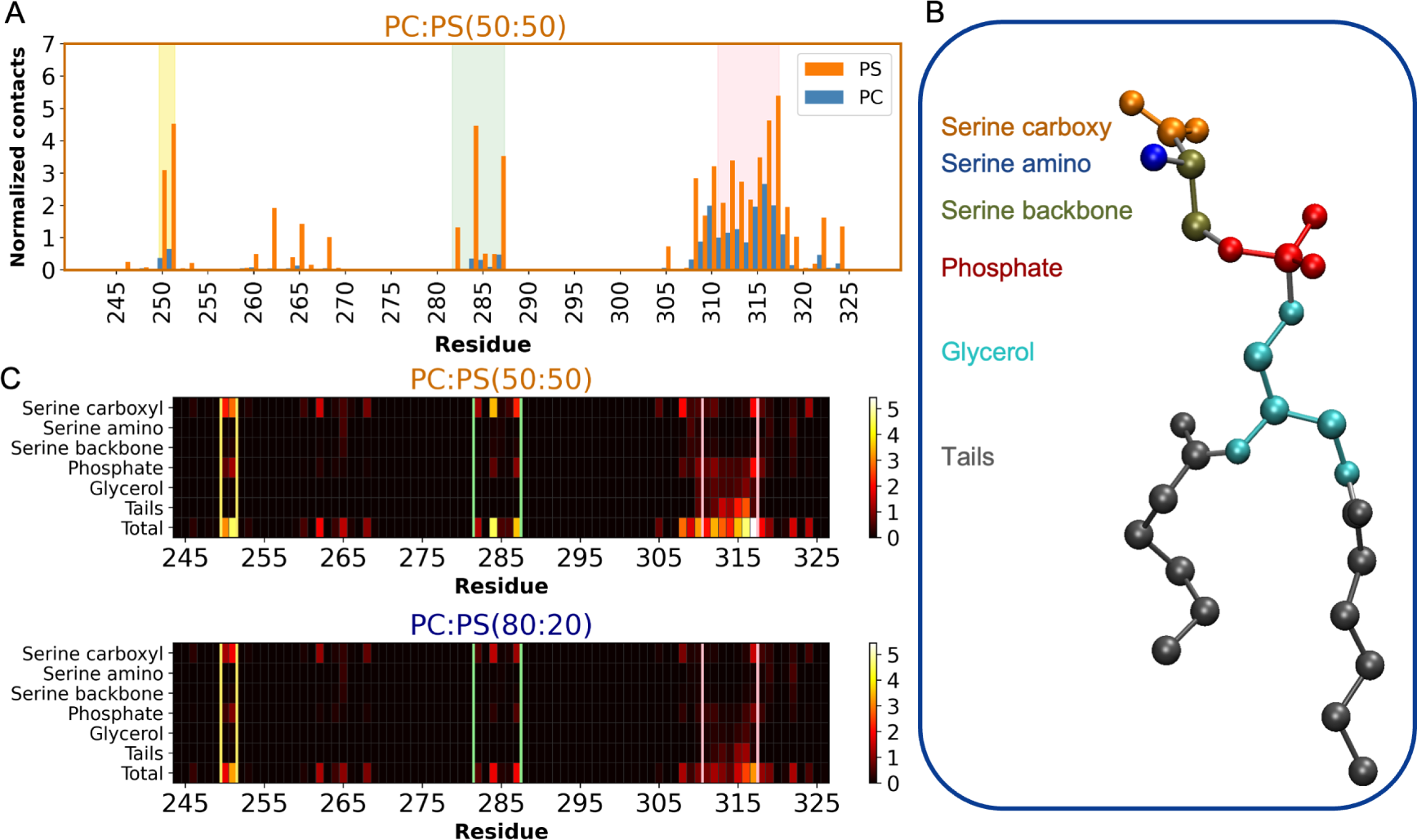
Preferential interaction of DV with anionic lipids. A) Contact map illustrating the number of interactions between DV residues and PC or PS lipids in the PC:PS (50:50) simulations. The total number of contacts for each residue and lipid type was calculated with Equation 1 (Fig. S4), and normalized by the total number of frames in which C*>*0. The lysine pair, the lysine-rich loop, and the hydrophobic loop are highlighted in yellow, green, and pink, respectively. PC and PS contacts are represented by blue and orange bars, respectively. B) Interaction with PS lipid heavy atoms broken into contributions from serine carboxy, serine amino, serine backbone, phosphate, glycerol, and tails, which are depicted in orange, blue, green, red, cyan, and gray, respectively. C) Heatmap showcasing the contact intensity between DV residues and PS lipid groups described in B. For PC:PS (50:50) and PC:PS (80:20) simulations, the total number of contacts for each residue and PS group was calculated with Equation 1 (Fig. S4), and normalized by the total number of frames in which C*>*0. Results for PC:PS (50:50) and PC:PS (80:20) are presented on top and bottom, respectively. The lysine pair, the lysine-rich loop, and the hydrophobic loop regions are denoted by yellow, green, and pink lines, respectively.

Collectively, our membrane-binding simulations substantiate earlier experimental evi-dence regarding the importance of surface-level, electrostatic interactions between the pos-itively charged lysine-rich loop and anionic lipids. They also corroborate the hypothesis of anchoring the hydrophobic loop into the membrane. Furthermore, our findings expand on previously well-studied membrane-binding hotspots (the lysine-rich loop and the hydropho-bic loop) by identifying the lysine pair as a novel membrane interaction site with comparable levels of lipid interaction to the lysine-rich loop.

### DV membrane binding modes

To investigate distinct membrane-binding modes of DV, in which protein-lipid interac-tions from various parts of DV lead to diverse orientations of *β*_2_GPI on the membrane sur-face, we first identified stable membrane-bound frames from our simulations through contact analysis and then grouped them into clusters. Specifically, we selected trajectory segments exhibiting stable “membrane-binding” events (see Methods for details) (Fig. S6). Out of 30 independent simulations with PS-containing membranes, 27 displayed stable membrane-bound DV configurations (Fig. S6A). For each membrane-bound configuration, we extracted the *z*-distance between the phosphate plane and individual DV residues (Fig. S6B) to dis-cern different binding modes of DV through clustering. Two distinct DV binding modes are illustrated in Fig. 4, Figs. S7 and S8. Binding mode 1 (cluster 1) includes 85% of the membrane-bound configurations, featuring the insertion of the hydrophobic loop into the membrane (Fig. 4A-B-D). Additionally, the positively charged lysine-rich loop and the ly-sine pair in this mode are in close proximity to the negatively charged lipid headgroups in the membrane (Fig. 4B-D). The anchoring of the hydrophobic loop to the membrane and favorable electrostatic interactions between positively charged lysine-rich regions and anionic lipids contribute to the stability of this binding mode, making it the dominant form observed in our simulations. A representative snapshot of binding mode 1 is shown in Fig. 4D, empha-sizing DV’s preference to bind to a PS-rich regions of the membrane, with its hydrophobic loop nestled between lipid tails, and the lysine-rich loop and the lysine pair oriented toward the negatively charged headgroups of PS lipids.

**Figure 4:**
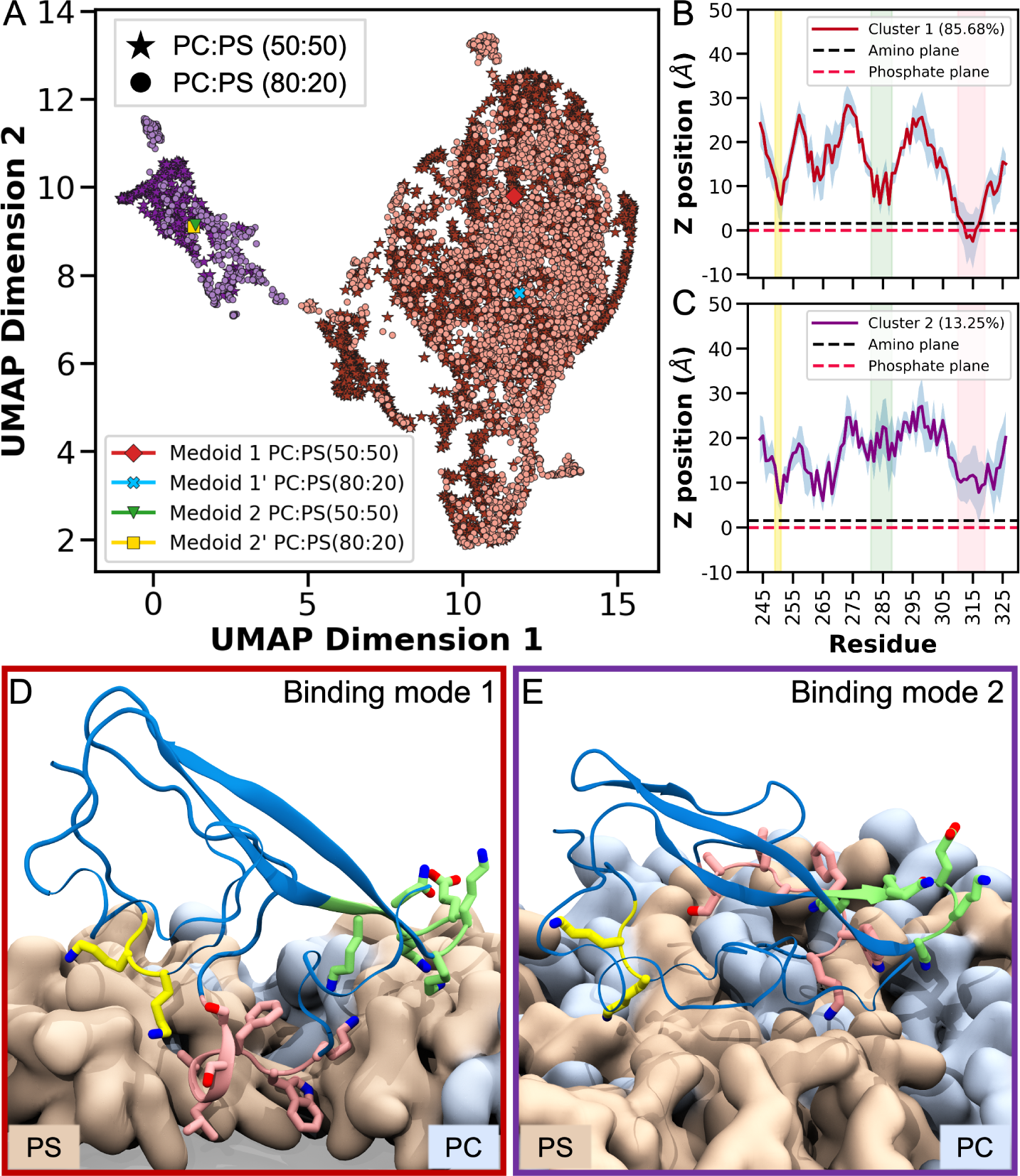
Clustering and characterization of DV membrane-bound poses. A) UMAP projection of membrane-bound DV poses, revealing distinct binding modes in clusters 1 (red) and 2 (purple). The results from PC:PS (50:50) and PC:PS (80:20) simulation data are represented by stars and circles, respectively. Medoids of cluster 1 are indicated by a red diamond (for PC:PS 50:50) and a blue cross (for PC:PS 80:20). Medoids of cluster 2 are denoted by a green triangle (for PC:PS 50:50) and a yellow square (for PC:PS 80:20). A medoid represents the membrane-bound pose with the lowest total pairwise distance to the cluster average (see Methods for details). B) *z*-distance between the side chain COM and the phosphate plane averaged for cluster 1 is drawn as a red line, with the standard deviation shown as a blue region. C) *z*-distance between the side chain COM and the phosphate plane averaged for cluster 2 is drawn as a purple line, with the standard deviation shown as a blue region. D) Medoid 1, illustrating membrane interactions in a representative snapshot for binding mode 1. E) Medoid 2, depicting superficial membrane interactions in a representative snapshot for binding mode 2. The legends in B and C include the percentages of the total membrane-bound configurations within each cluster: 85.68% and 13.25%, respectively, for binding modes 1 and 2. The position of phosphate and amino planes are indicated as red and black dashed lines, respectively. The lysine pair, the lysine-rich loop, and the hydrophobic loop regions are highlighted in yellow, green, and pink, respectively. In D and E, DV is presented in blue, with the the lysine pair, the lysine-rich loop, and the hydrophobic loop highlighted in yellow, green, and pink, respectively. PC and PS lipids are shown in blue and beige, respectively.

Binding mode 2 (cluster 2), which is observed only for 13% of the membrane-bound DV configurations, can be characterized as a more superficial DV positioning on top of the membrane surface without any clear insertion (Fig. 4A-C-E). We believe that binding mode 2 only represents an intermediate state for DV binding to the membrane arising in our simulations, and not a stable binding state. A representative snapshot of binding mode 2 is shown in Fig. 4E, illustrating how DV might approach a PS-rich section of the membrane, with the lysine pair and lysine-rich loop residues directed towards the anionic lipid head groups. Taken together, our findings indicate that the membrane-bound state of DV (binding mode 1), captured in the majority of the simulations, is stabilized by the insertion of its hydrophobic loop into the membrane and attractive electrostatic interactions of its two positively charged regions with anionic phospholipids.

To better characterize DV orientation with respect to the membrane in different membrane binding modes, we quantified the angle between the three principal axes (PAs) of DV and the membrane normal (Fig. 5). In binding mode 1, all three angles exhibit narrow distributions, suggesting a more stable binding to the membrane. Conversely, in binding mode 2, the PA1 angle shows a narrow distribution around 97.1*^◦^*, but PA2 and PA3 angles span a broad range, indicating a wide range of motion atop the membrane and a lack of stabilization, possibly attributed to the absence of hydrophobic loop anchoring (Fig. 4D-E).

**Figure 5:**
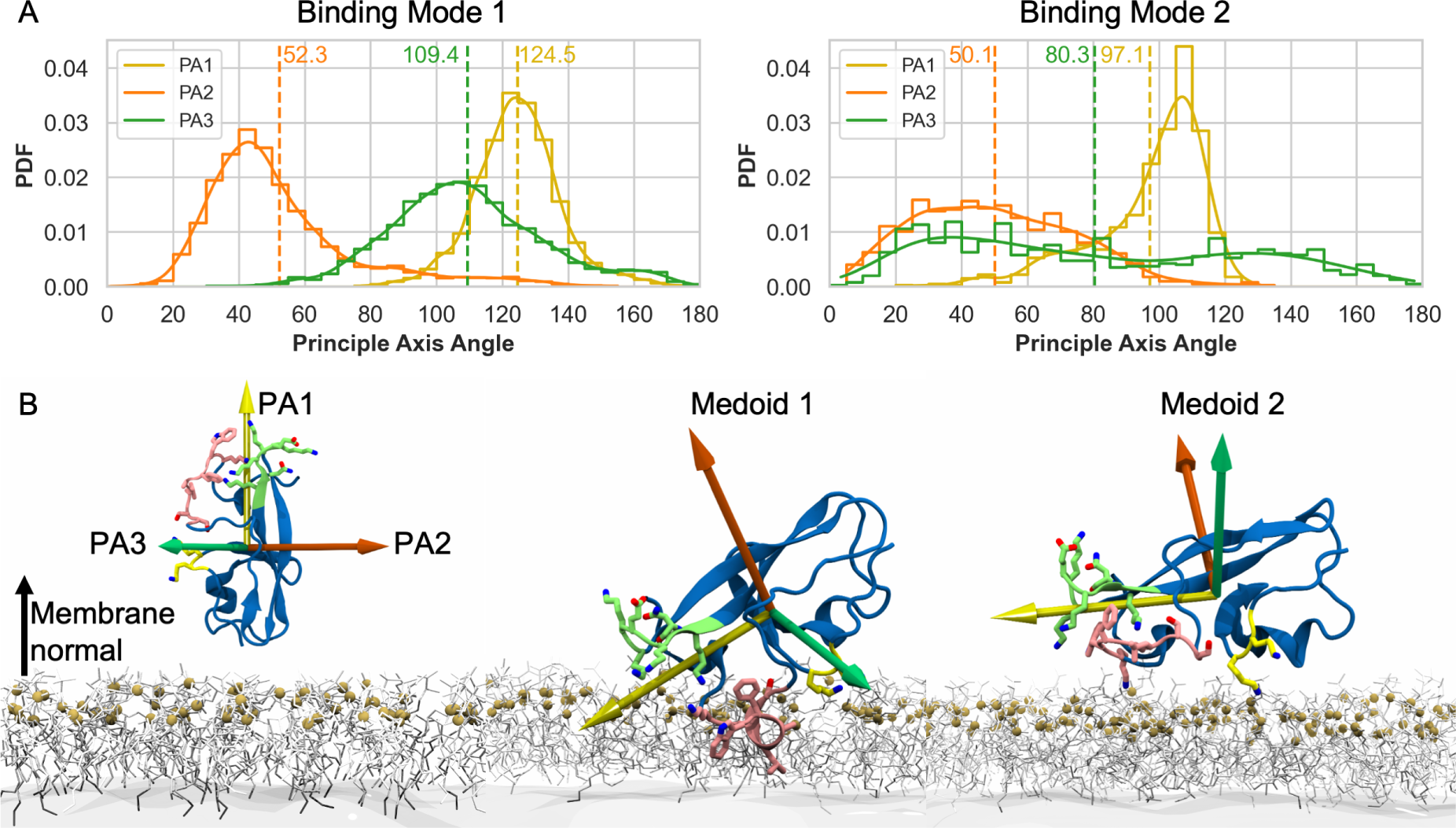
Orientation of membrane-bound DV. A) Probability density function (PDF) illustrating the angle between DV’s three principal axes (PA1 in yellow, PA2 in orange, and PA3 in green) and the membrane normal in two binding modes (clusters). Results for binding mode 1 are presented on the left, and for binding mode 2 on the right. The angle distribution, sampled from the simulations, is presented both as normalized histograms (depicted as stairstep plots) and as PDFs (represented by solid lines). These PDFs are calculated using Gaussian kernel density estimation applied to the histograms. Vertical dashed lines, labeled with their corresponding colors, indicate the average angle for each PA. B) From left to right, the orientation of principal axes (PA1, PA2, and PA3) for the initial placement of DV, for binding mode 1, and for binding mode 2, respectively. DV is drawn in blue, with the lysine pair, the lysine-rich loop, and the hydrophobic loop highlighted in yellow, green, and pink, respectively. Phospholipids are shown in gray, and the organic solvent in the membrane core is represented as gray surface. The phosphorous atoms of lipids are highlighted as tan-colored spheres.

To assess the stability of the dominant binding mode (binding mode 1) identified in our HMMM simulations, we converted a representative frame (medoid 1) from HMMM to full membrane and conducted an additional 300-ns simulation (Fig. 6). During the extended, full-membrane simulation, the hydrophobic loop remained inserted into the membrane.

**Figure 6:**
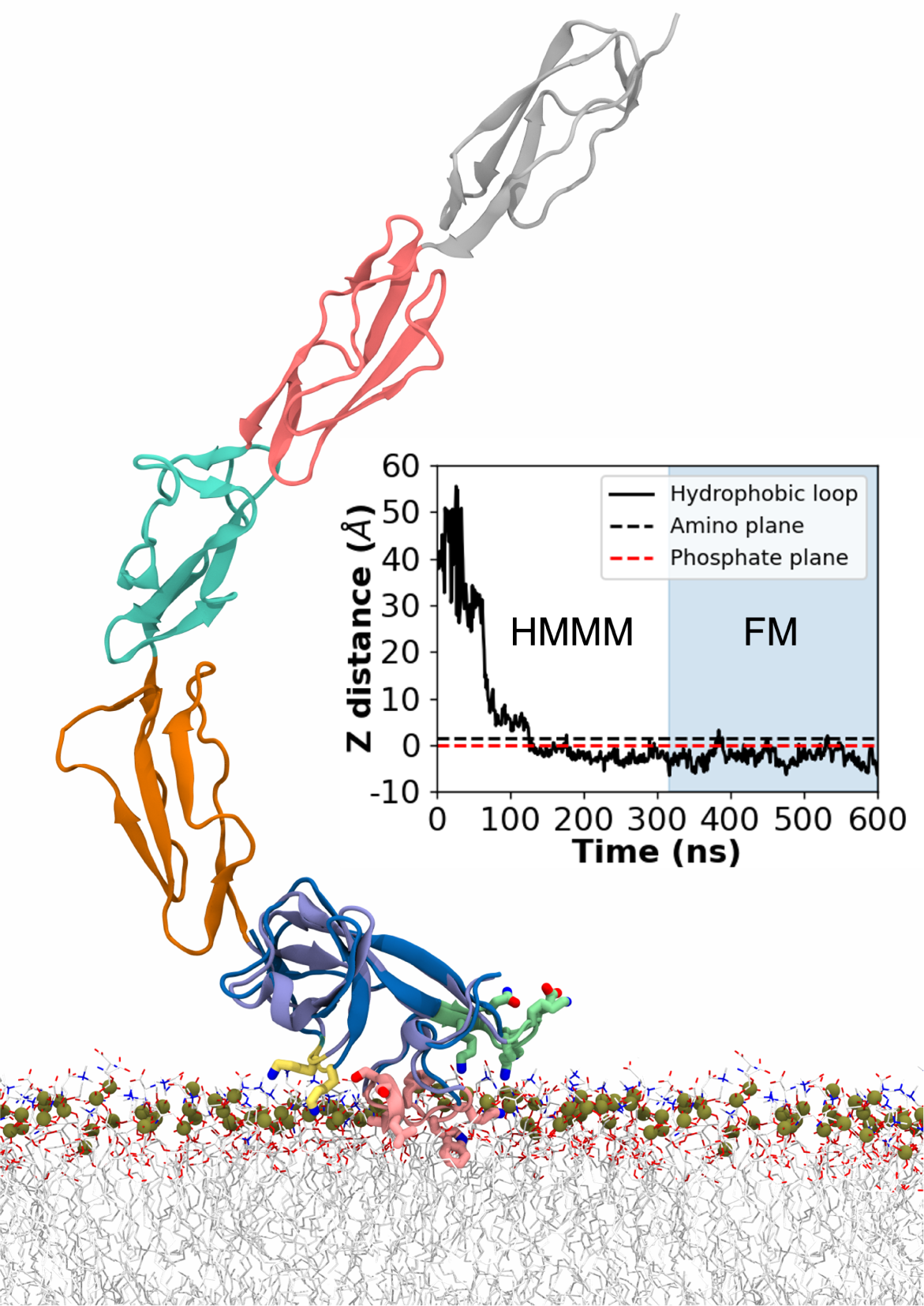
Superposition of *β*_2_GPI on full membrane-bound DV following conversion from HMMM to full membrane (FM) An X-ray crystal structure displaying the J-elongated confor-mation of *β*_2_GPI is overlaid on the full membrane-bound DV. Domains DI to DV are represented in gray, pink, green, orange, and light blue, respectively. The membrane-bound DV is depicted in dark blue, with the lysine pair, the lysine-rich loop, and the hydrophobic loop high-lighted in yellow, green, and pink, respectively. The lipid phosphorous atoms are drawn as tan-colored spheres. The inset illustrates the time evolution of *z*-distance between the side chain COM of the hydrophobic loop and the phosphate plane, spanning the initial 300 ns of HMMM membrane binding and the subsequent 300 ns of full membrane sim-ulations (highlighted in a blue background) for binding mode 1. The phosphate and amino planes are shown as red and black dashed lines, respectively.

Finally, in order to assess the compatibility of this binding mode to membrane binding of the whole protein, the full *β*_2_GPI structure was superimposed onto the DV. The resulting structure showed that the binding mode captured by the MD simulations does indeed allow for full *β*_2_GPI in its J-elongated conformation to be on the membrane (Fig. 6) without any steric clashes. This observation suggests that the full membrane-bound DV is compatible with the antigenic function of *β*_2_GPI in APS, while keeping aPLs epitopes on DI accessible (Fig. 6). These findings support the robustness and relevance of the captured membrane-bound DV in a more physiological context.

## Conclusion

*β*_2_GPI is a major protein antigen in the autoimmune thrombotic disorder known as APS, which is recognized for its significant morbidity rate, notably as a leading cause of stroke in individuals younger than 50. ^93^ A crucial aspect of this disease is the interaction of *β*_2_GPI with anionic phospholipids, which makes it a more effective antigen for antiphospholid antibodies. However, it is still unclear how *β*_2_GPI interacts with anionic phospholipids and acquires this pathogenic function at ana tomic level. This study is the first to explore the binding of *β*_2_GPI to anionic membranes at an atomic level, filling this knowledge gap.

It is recognized that DV in *β*_2_GPI plays a central role in its membrane binding. Despite insights from previous experimental studies, the details of interactions between DV and membrane remained poorly understood. Here, through a large set of atomistic simulations, we provide the first atomic structure of membrane-bound DV and how anionic lipids are important for its binding. Membrane association of *β*_2_GPI DV is primarily facilitated via electrostatic interactions of lysine-rich regions with the head groups of anionic lipids and insertion of a hydrophobic loop into the core of the membrane.

MD simulations have been successfully used to characterize similar peripheral protein systems but never to study *β*_2_GPI interaction with an-ionic lipids.

Previous biochemical studies had provided evidence for critical requirement of the lysine-rich loop (K282-K287) and the hydrophobic loop (S311-K317) in DV for membrane binding. Our simulations clearly show at an atomic level how these parts of the protein participate in membrane binding.

Furthermore, we have dis-covered that the binding inter-face of DV is more extended than was previously thought (Figs. 2 and 3 and Figs. S1 to S5). The results reveal that the lysine pair (K250/K251) also strongly contributes to binding, further stabilizing the interaction of *β*_2_GPI with the anionic lipid surface. Together, these findings offer a structural framework for rationally targeting DV to block its lipid binding.

Next, we identified two distinct membrane-bound configurations for DV of *β*_2_GPI, which differ primarily with regard to the extent of the hydrophobic loop insertion into the membrane (Fig. 4). In mode 1, the hydrophobic loop is fully inserted, but not in mode 2. Given the energetically favorable contacts between hydrophobic W316 and the lipid tails, mode 1 is found to be the dominant mode in our simulations. This strongly indicates that mode 1 is the primary configuration for DV’s membrane-bound state. We believe this mode represents the physiological membrane binding mode of DV. However, the significance of mode 2 should not be downplayed, as this mode could be an intermediate for membrane binding or relevant under physiological situations where *β*_2_GPI simultaneously interacts with other negatively charged surfaces like heparin and polyphosphates, both of which are known to regulate many aspects of the coagulation cascade.

Finally, in this study, we focused on the oxidized form *β*_2_GPI, which is characterized by an intact C-terminal disulfide bond (C288-C326). However, a reduced form of *β*_2_GPI exists *in vivo*, too.^13,94^ Unlike the oxidized form, the reduced form, which is characterized by the broken disulfide bond (C288-C326), is incapable of binding to the membranes. ^22^ Given that a U-shaped conformation of the hydrophobic loop in mode 1 is crucial in anchoring DV to the lipids, it is tempting to speculate that the structural basis behind the loss of phospholipid binding in reduced *β*_2_GPI is the linearization of this loop. Moving forward, investigating the distinctions between the oxidized and reduced forms of *β*_2_GPI will be essential to test this hypothesis, potentially uncovering new regulatory mechanisms that affect the membrane interaction of *β*_2_GPI. Mutagenesis experiments guided by these new structural models could also be pursued to enrich our understanding of *β*_2_GPI-lipid interactions. Together, this work could lead to the identification of novel therapeutic approaches to block the binding of *β*_2_GPI to the membranes, which would be transformative for APS patients.

## Acknowledgement

Research reported in this publication was supported by the National Institutes of Health under award numbers P41-GM104601, R24-GM145965 and R01-GM123455 (to ET) and R01-HL150146 (to NP). We also acknowledge computing resources provided by NSF ACCESS (MCA06N060 to ET) at National Supercomputing Centers.

## Author Contributions

H.S.H., N.P. and E.T. designed the research. H.S.H carried out all simulations and analyzed the data. All authors wrote the article.

