## Supplementary Information for "Atomistic Characterization of Beta-2-Glycoprotein I Domain V Interaction with Anionic Membranes"

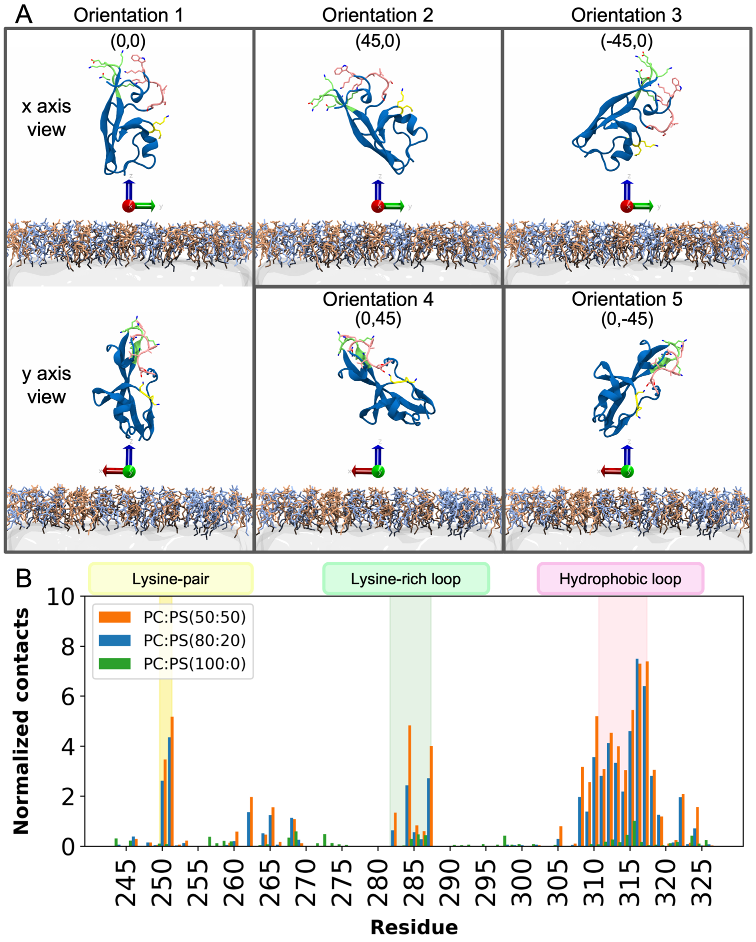


Figure S1: **Initial orientations of DV in membrane binding simulations and contact map between DV residues and the membrane with different lipid compositions.** A) Two panels on the left depicts orientation 1, serving as the reference orientation with no rotation along the $x$ and $y$ axes (indicated as (0,0)). The top middle and top right panels display orientations 2 and 3, with DV rotated by 45 and $-45$ degrees along its $x$-axis, respectively. The bottom middle and bottom right panels show orientations 4 and 5, with DV rotated by 45 and $-45$ degrees along its $y$-axis, respectively. DV is illustrated in dark blue, while the lysine pair, the lysine-rich loop, and the hydrophobic loop are highlighted in yellow, green, and pink, respectively. PC and PS lipids are differentiated in blue and orange, respectively. Reference axes $(x, y, z)$ are denoted in red, green, and blue beneath DV. B) The total number of contacts with the membrane for each residue, calculated using Equation 1 and normalized based on the number of frames with C>0 for each lipid composition. The regions corresponding to lysine pair, the lysine-rich loop, and the hydrophobic loop are highlighted in yellow, green, and pink, respectively. Results for membrane compositions PC:PS (50:50), PC:PS (80:20), and PC:PS (100:0) are represented by orange, blue, and green bars, respectively.

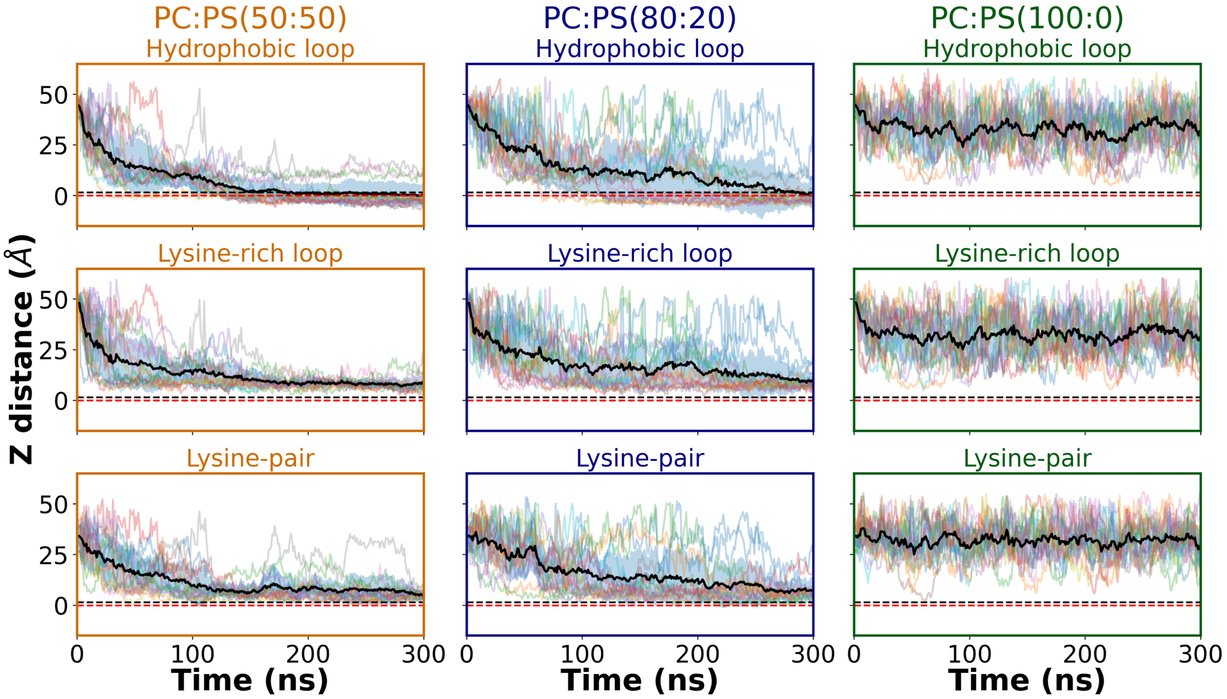


Figure S2: **Spatiotemporal dynamics of DV membrane interaction hotspots during membrane binding.** Time evolution of $z$-distance between the phosphate plane and the side chain COM averages for the hydrophobic loop, the lysine-rich loop, and the lysine pair. Results for membrane compositions PC:PS (50:50), PC:PS (80:20) and PC:PS (100:0) are presented in orange, blue and green panels, respectively. Lighter colored lines represent $z$-distance between the side chain COM of the selected region and the phosphate plane for individual replicas. The average $z$-distance of all replicas is depicted in black, with the standard deviation shown as a blue shadow. The positions of phosphate and amino average planes are indicated by red and black dashed line, respectively.

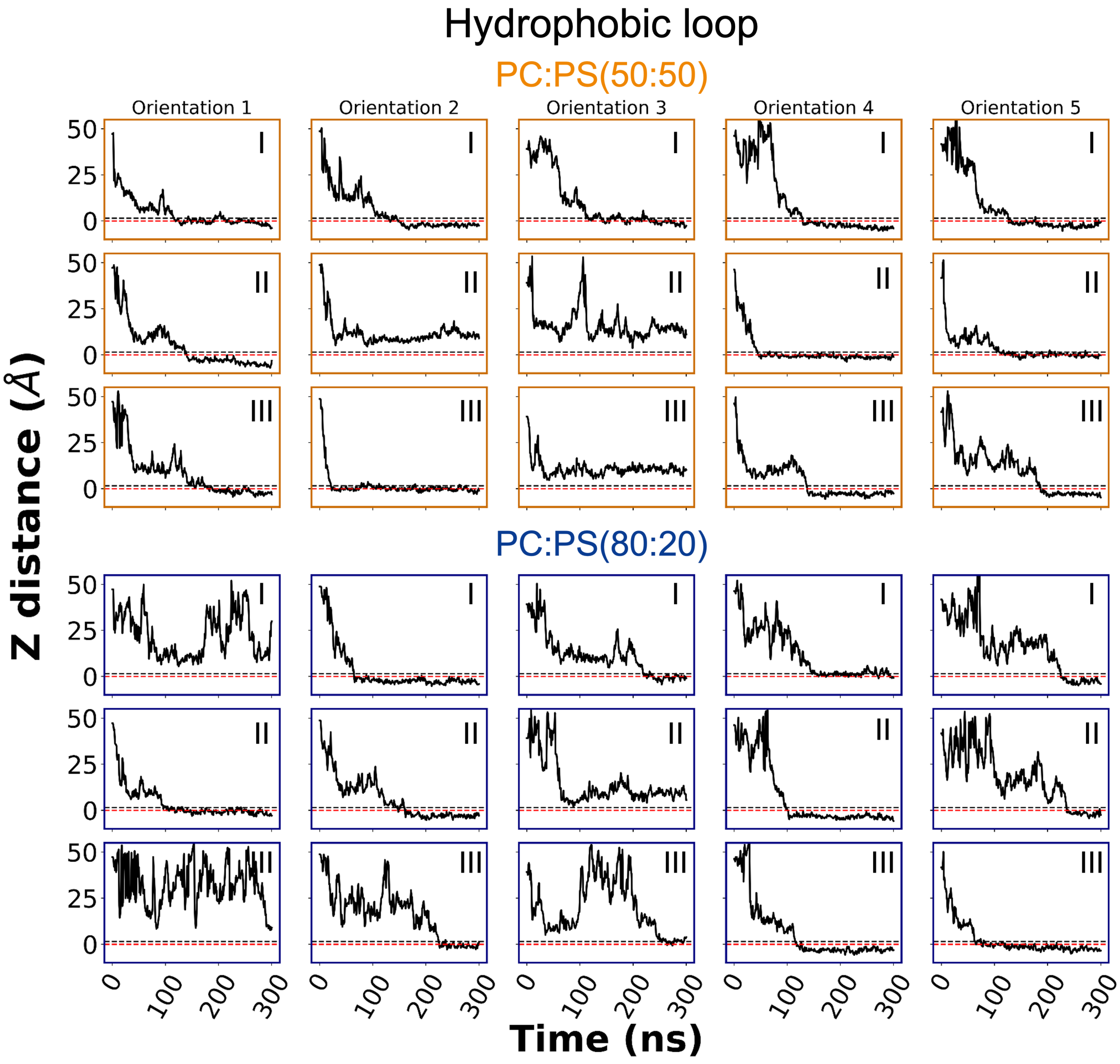


Figure S3: **Spontaneous insertion of the hydrophobic loop under the phosphate plane during 30 independent membrane-binding simulations.** Time evolution of $z$-distance between the side chain COM average for the hydrophobic loop and the phosphate plane. Results for membrane compositions PC:PS (50:50) and PC:PS (80:20) are presented in orange and blue panels, respectively. Each column displays plots for three replicas (replica number indicated in the upper right corner) sharing the same initial protein orientation as defined in (Fig. S1). The positions of phosphate and amino average planes are indicated by red and black dashed line, respectively.

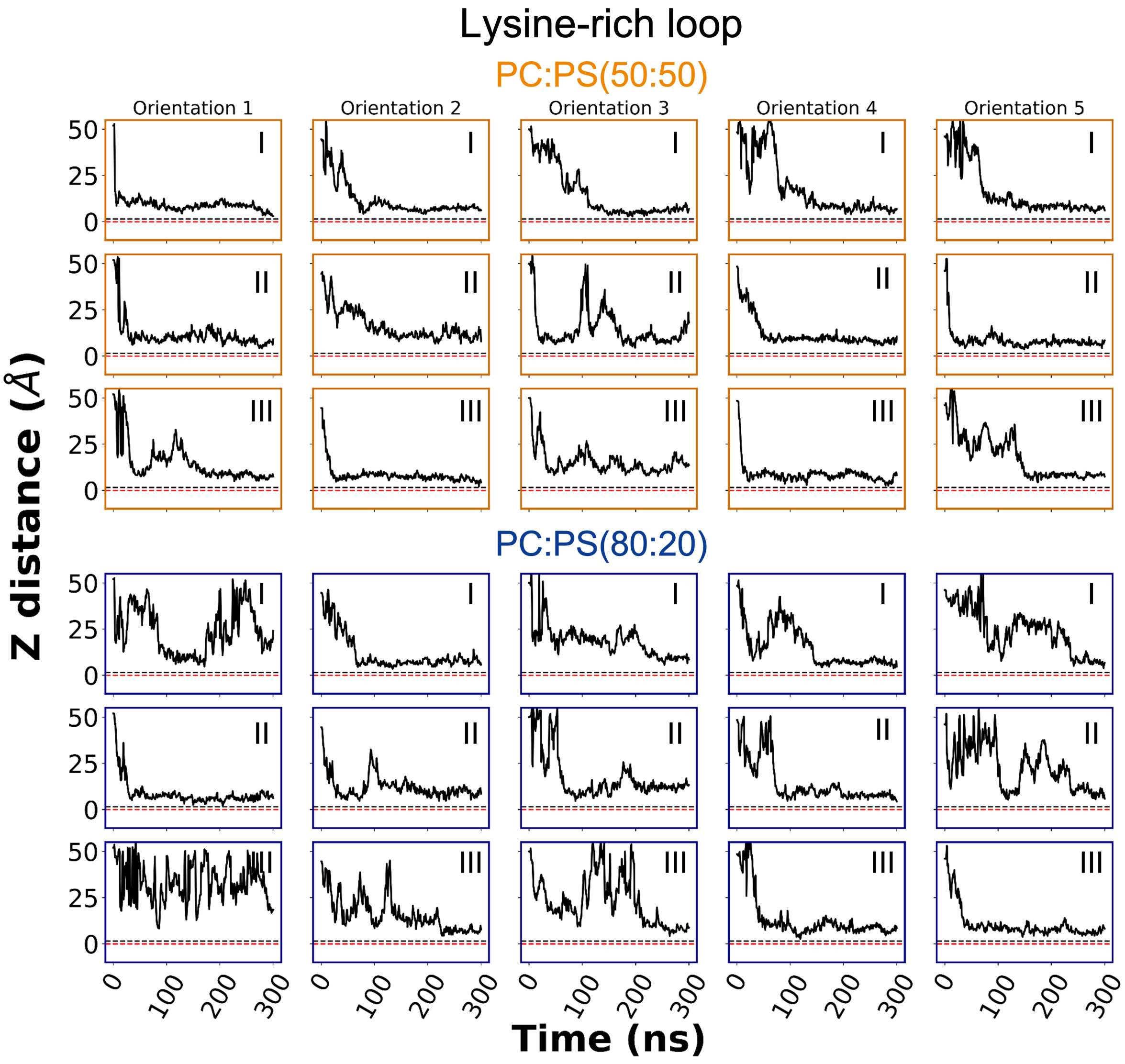


Figure S4: **Surface-level interactions of the lysine-rich loop with the membrane in 30 independent membrane-binding simulations.** Time evolution of $z$-distance between the side chain COM average for the lysine-rich loop and the phosphate plane. Results for membrane compositions PC:PS (50:50) and PC:PS (80:20) are presented in orange and blue panels, respectively. Each column displays plots for three replicas (replica number indicated in the upper right corner) sharing the same initial protein orientation as defined in (Fig. S1). The positions of phosphate and amino average planes are indicated by red and black dashed line, respectively.

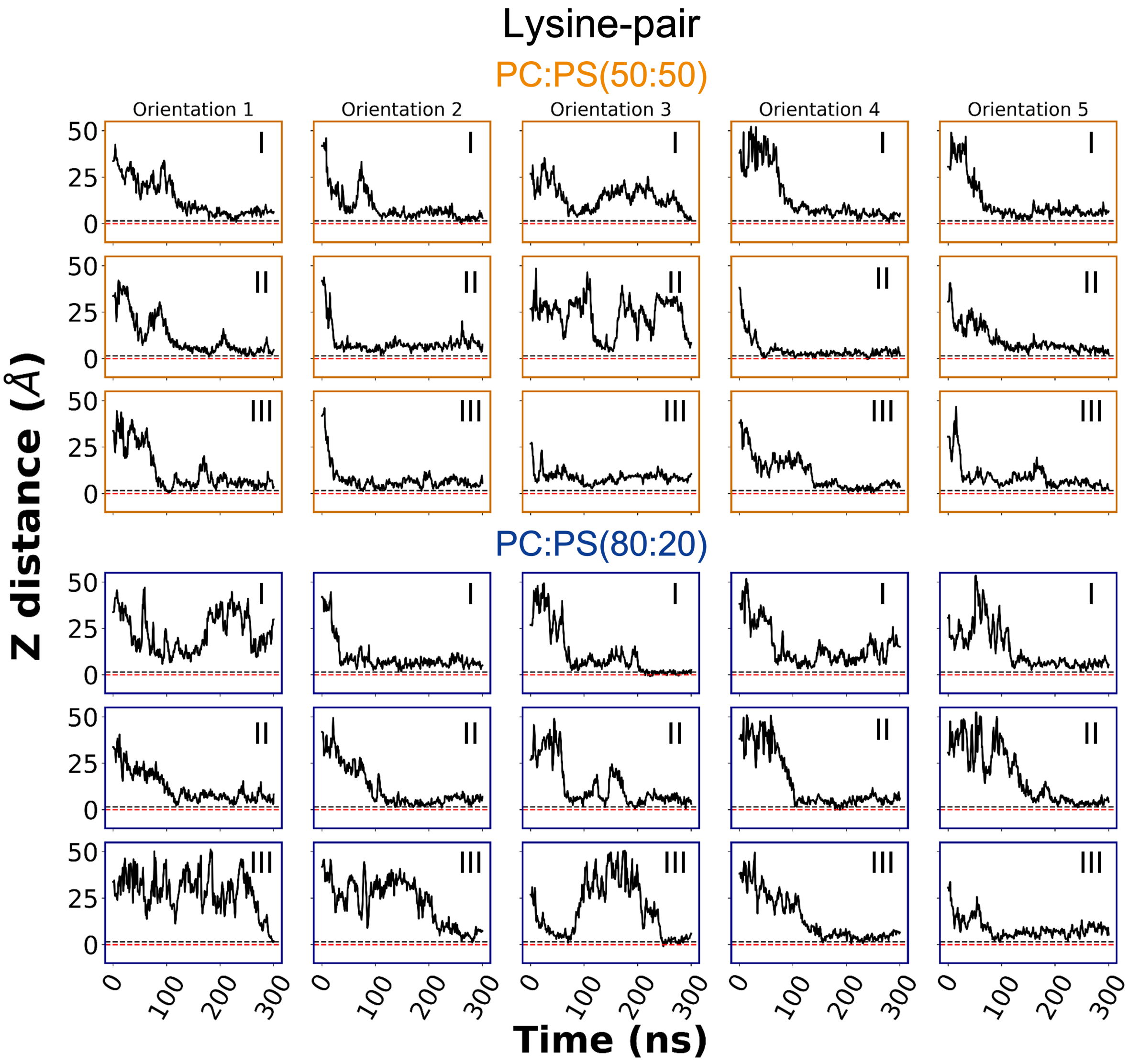


Figure S5: **Surface-level interactions of the lysine-pair with the membrane during 30 independent membrane-binding simulations.** Time evolution of $z$-distance between the side chain COM average for the lysine-pair and the phosphate plane. Results for membrane compositions PC:PS (50:50) and PC:PS (80:20) are presented in orange and blue panels, respectively. Each column displays plots for three replicas (replica number indicated in the upper right corner) sharing the same initial protein orientation as defined in (Fig. S1). The positions of phosphate and amino average planes are indicated by red and black dashed line, respectively.

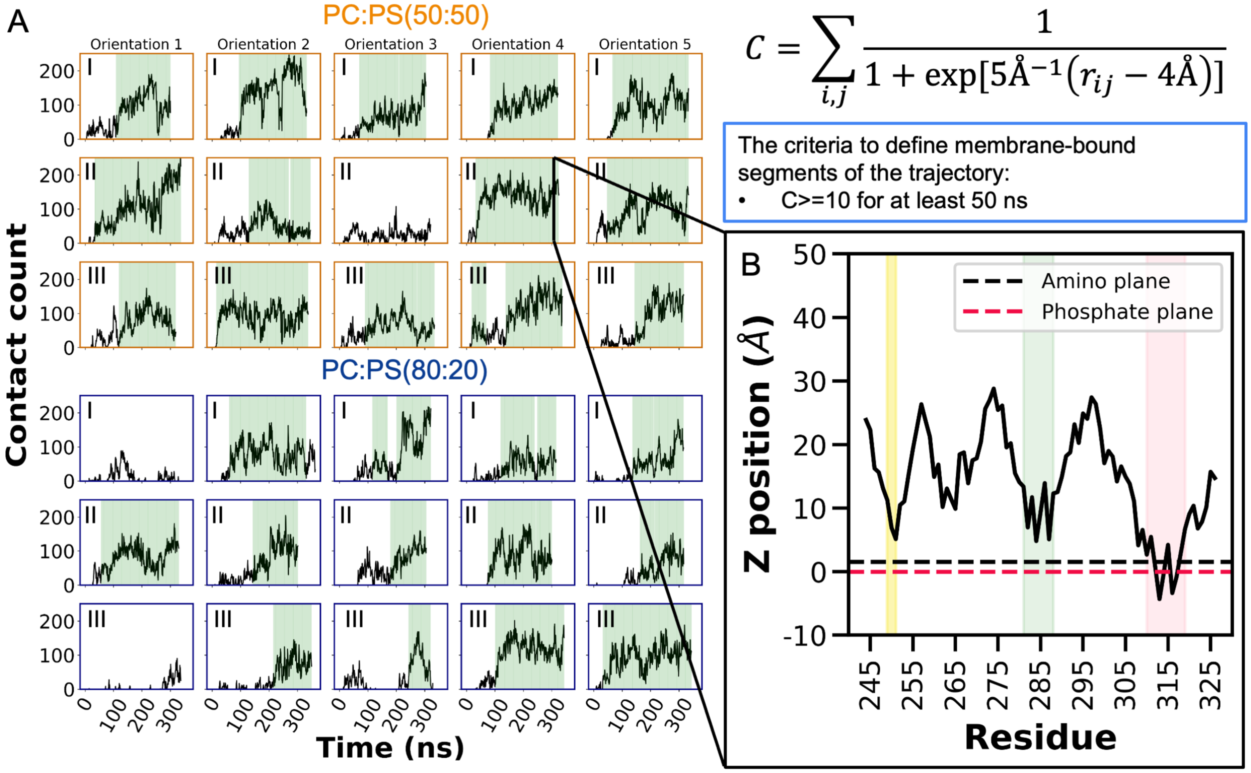


Figure S6: **Defining the trajectory segments with stable membrane-bound configurations based on the number of membrane-DV contacts.** A) The figure illustrates the total number of contacts between DV and the membrane, calculated using Equation 1 (also shown on the top right), at 1-ns intervals. Results for membrane compositions PC:PS (50:50) and PC:PS (80:20) are presented in orange and blue panels, respectively. Each column displays plots for three replicas (replica number indicated in the upper left corner) sharing the same initial protein orientation as defined in (Fig. S1). The segments of the trajectories in which stable binding (criteria summarized in the blue box, see Methods for details) was observed are marked with a green background. B) A representative snapshot displays the distance between the side chain COM and the phosphate plane of a membrane-bound frame. The positions of phosphate and amino average planes are indicated by red and black dashed line, respectively. The lysine pair, the lysine-rich loop, and the hydrophobic loop regions are highlighted in yellow, green, and pink, respectively.

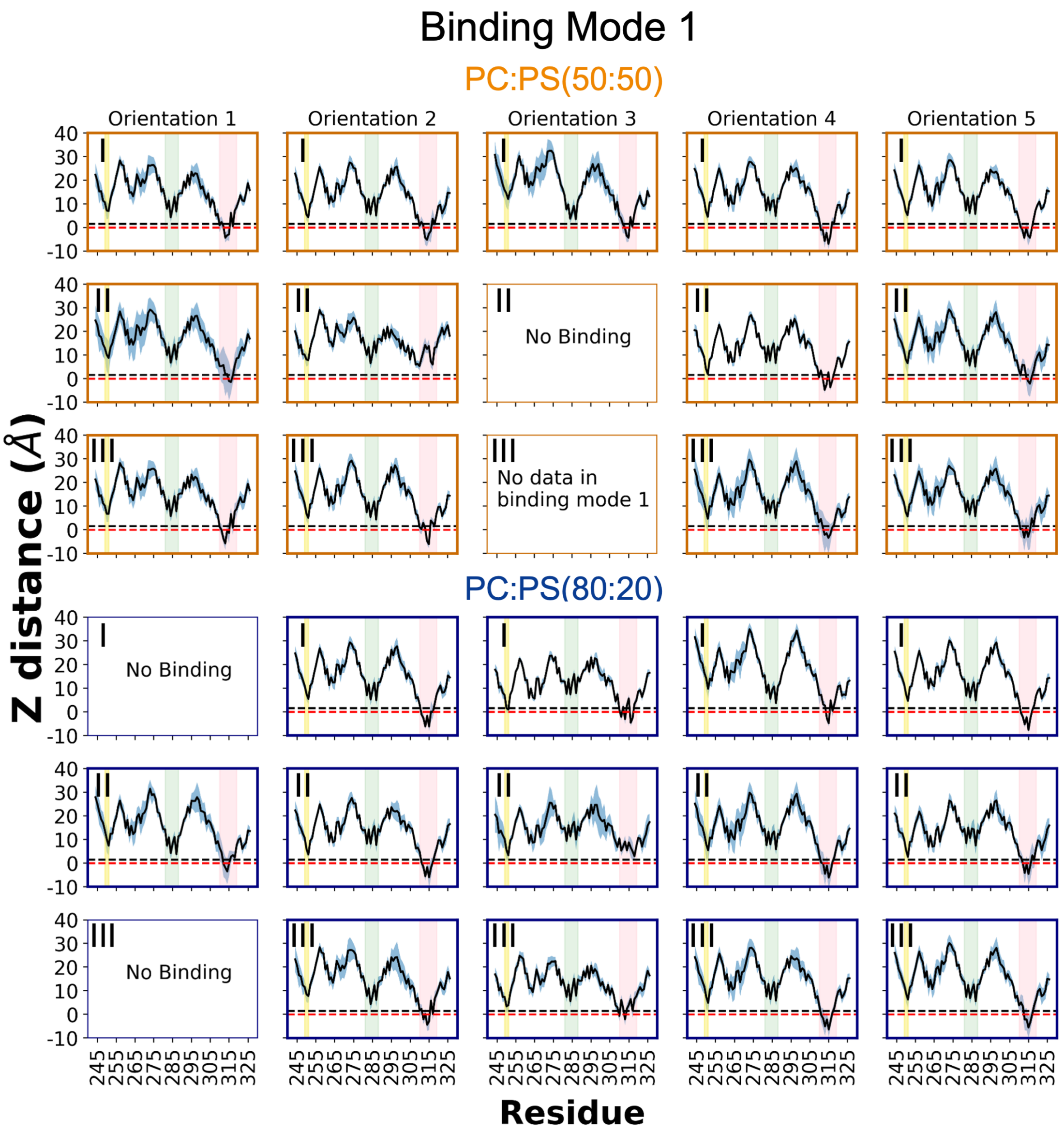


Figure S7: **Average position of each residue with respect to the membrane in binding mode 1 across 30 independent membrane-binding simulations.** Ensemble average of $z$-distance between the side chain COM and the phosphate plane from the membrane-bound frames in binding mode 1 cluster, with the standard deviation shown as a blue shadow. Results for membrane compositions PC:PS (50:50) and PC:PS (80:20) are presented in orange and blue panels, respectively. Each column displays plots for three replicas (replica number indicated in the upper right corner) sharing the same initial protein orientation as defined in (Fig. S1). The positions of phosphate and amino average planes are indicated by red and black dashed line, respectively. The lysine pair, the lysine-rich loop, and the hydrophobic loop regions are highlighted in yellow, green, and pink, respectively.

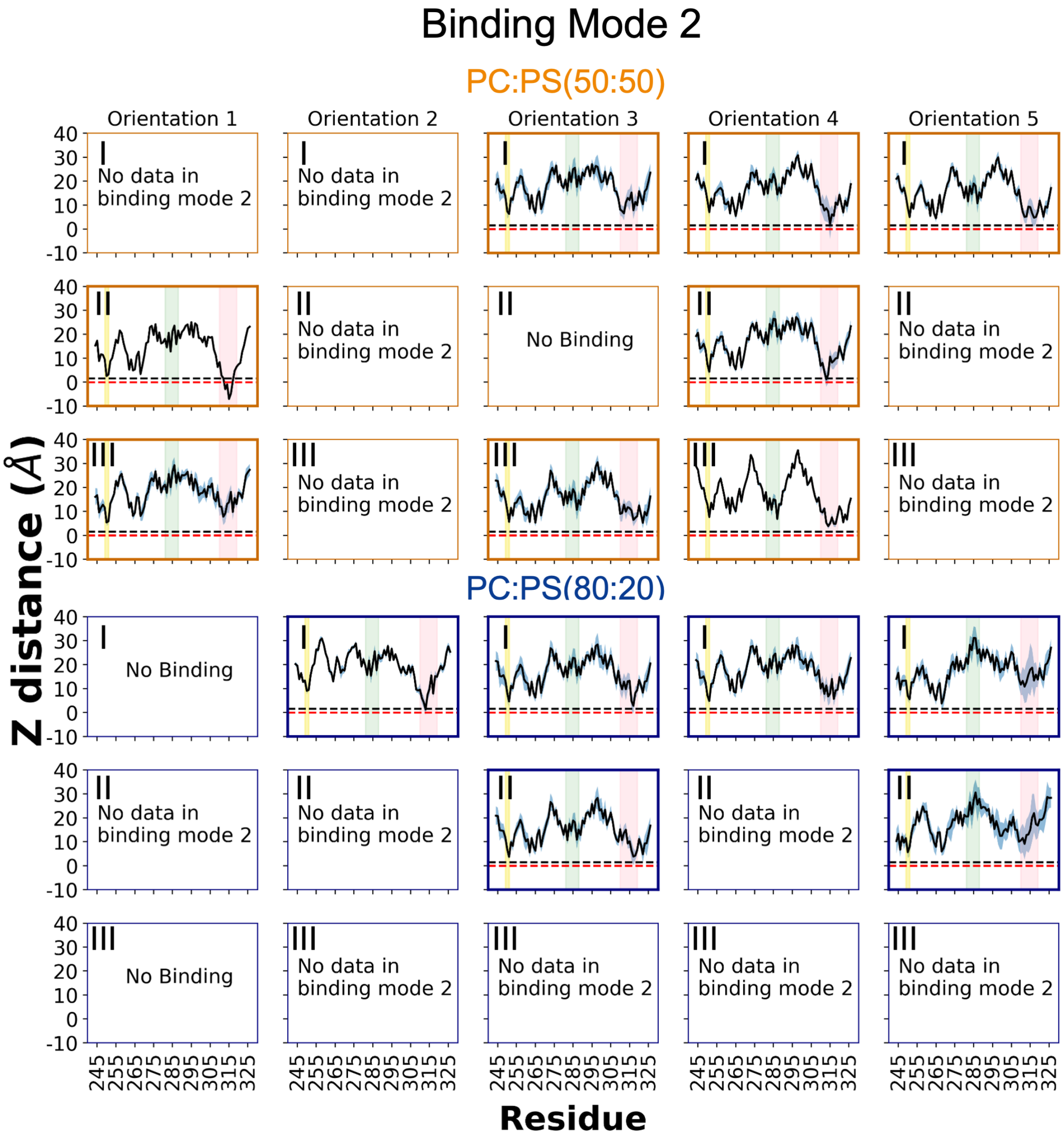


Figure S8: **Average position of each residue with respect to the membrane in binding mode 2 across 30 independent membrane-binding simulations.** Ensemble average of $z$-distance between the side chain COM and the phosphate plane from the membrane-bound frames in binding mode 2 cluster, with the standard deviation shown as a blue-colored region. Results for membrane compositions PC:PS (50:50) and PC:PS (80:20) are presented in orange and blue panels, respectively. Each column displays plots for three replicas (replica number indicated in the upper right corner) sharing the same initial protein orientation as defined in (Fig. S1). The positions of phosphate and amino average planes are indicated by red and black dashed line, respectively. The lysine pair, the lysine-rich loop, and the hydrophobic loop regions are highlighted in yellow, green, and pink, respectively.
